# Polymorphisms in the vitamin D system and mortality - The Tromsø study

**DOI:** 10.1101/588871

**Authors:** Rolf Jorde, Tom Wilsgaard, Guri Grimnes

## Abstract

**Background and objective:** Vitamin D deficiency is associated with diabetes, cancer, immunological and cardiovascular diseases as well as increased mortality. It has, however, been difficult to show a causal relation in randomized, controlled trials. Mendelian randomization studies provide another option for testing causality, and results indicate relations between the serum 25-hydroxyvitamin D (25(OH)D) level and some diseases, including mortality. We have from the Tromsø Study in 2012 published non-significant relations been vitamin D related single nucleotide polymorphisms (SNPs) and mortality, but have since then genotyped additional subjects, the observation time is longer and new SNPs have been included.

**Methods:** Genotyping was performed for SNPs in the *NADSYN1*, *CYP2R1*, *VDR, CUBILIN* and *MEGALIN* genes in 11 897 subjects who participated in the fourth survey of the Tromsø Study in 1994-1995. Serum 25(OH)D levels were measured in 6733 of these subjects. A genotype score based on SNPs in the *NADSYN1* and *CYP2R1* genes (related to the serum 25(OH)D level) and serum 25(OH)D percentile groups were created. Mortality data was updated till end of March 2017 and survival analysed with Cox regression adjusted for sex and age.

**Results:** During the observation period 5491 subjects died. The genotype score and the serum 25(OH)D percentile groups were (without Bonferroni correction) significantly related to mortality in favour of high serum 25(OH)D. None of the SNPs in the *VDR* or *MEGALIN* genes were related to mortality. However, for the rs12766939 in the *CUBILIN* gene with the major homozygote as reference, the hazard ratio for mortality for the minor homozygote genotype was 1.17 (1.06 – 1.29), P < 0.002. This should be viewed with caution, as rs12766939 was not in Hardy-Weinberg equilibrium.

**Conclusion:** Our study confirms a probable causal but weak relation between serum 25(OH)D level and mortality. The relation between rs12766939 and mortality needs confirmation in more homogenous cohorts.

## Introduction

The nuclear vitamin D receptor (VDR) is found in most tissues of the body, and enzymes necessary for the activation of vitamin D to 25-hydroxyvitamin D (25(OH)D), and finally to 1,25-dihydroxyvitamin D (1,25(OH)_2_D), are located not only in the liver and kidneys, but in peripheral tissues as well. Vitamin D is essential for calcium absorption and bone health, but may also have a number of other biological functions, in particular related to cell proliferation and immunology [1].

Low serum levels of 25(OH)D, which is used as a marker of the body’s vitamin D stores, are associated with cardiovascular risk factors like hypertension, hyperglycaemia and hyperlipidaemia, as well as manifest diseases like cancer, type 2 diabetes, cardiovascular and immunological diseases [2]. It has, however, been difficult to show a beneficial effect of vitamin D supplementation in treatment or prevention of these diseases in randomized, controlled trials (RCTs), possibly because most of those included have not been in need of supplementation as their 25(OH)D status has been more than adequate [3,4]. There are also ethical problems with including vitamin D deficient subjects for an extended period of time, and one may therefore not get the final answer regarding vitamin D supplementation from RCTs.

Another approach is the Mendelian randomization (MR) procedure [5], and there are several single nucleotide polymorphisms (SNPs) related to enzymes needed for activation, transport and breakdown of vitamin D, as well as in the VDR. However, the results have not consistently been in favour of vitamin D. Thus, SNPs related to synthesis and breakdown of 25(OH)D have been associated with type I diabetes [6], hypertension [7] and multiple sclerosis [8], whereas not to cardiovascular disease [9], fractures [10], or type 2 diabetes [11]. Similarly, there have been numerous studies on *VDR* SNPs, in particular regarding cancer, but also these have shown conflicting results [12,13].

One explanation could be that the effect of vitamin D is modest (if present at all), and therefore a large number of subjects are needed to show an effect on specific diseases. On the other hand, if there is a broad-ranging effect of vitamin D as indicated by the overwhelmingly positive effect in observational studies and the wide-spread localization of the VDR, one would assume that this would add up to an increased mortality risk in those with vitamin D deficiency that would be more easy to demonstrate [14].

A relation to mortality has also been found in meta-analyses of vitamin D RCTs [15,16], but none of the included studies were specifically designed for that purpose, and confirmation in more trials is needed. Similarly, in a large MR study using SNPs associated to the serum 25(OH)D level, genetically low 25(OH)D was related to mortality [17], but again, more studies were asked for [18]. There are also a few MR studies on mortality using *VDR* SNPs, but these have been too small to draw firm conclusions [19–21].

We have previously reported a lack of significant association between 25(OH)D related SNPs and mortality in 9528 subjects followed for up to 15 years in the Tromsø study [22]. Since then we have genotyped additional subjects, included genotyping of *VDR*, *MEGALIN* and *CUBILIN* SNPs, and the observation period is now up to 22 years. In view of the uncertainty regarding vitamin D and mortality, we therefore found it worthwhile to reanalyse the cohort for vitamin D SNPs and mortality.

## Methods

### Subjects

The Tromsø study is a repeated population-based study conducted in the municipality of Tromsø, Norway, situated at 69°N (current population 76 000). The study was initiated in 1974, and has been performed seven times at regular intervals. The seventh and latest survey was conducted in 2015-2016. In the fourth survey in 1994-1995, all individuals aged 25 years or older and living in Tromsø were invited. A total number of 27 158 persons participated in the first visit, providing an attendance rate of 77%. All men aged 55–74 years, all women aged 50–74 years and a sample of 5–10% of the remaining age groups between 25 and 84 years were invited to a second visit with more extensive clinical examination, and 7965 persons, or 78% of those invited, attended [23]. The study was conducted by UiT The Arctic University of Norway in cooperation with the National Health Screening Service. For this cohort of 27 158 subjects, specific endpoint registers for myocardial infarction, type 1 diabetes, stroke, hip and radial fractures, and aortic stenosis have been created. In addition, data from the Cancer Registry of Norway and the National Causes of Death Registry were available. In our initial report in 2012, subjects with one or more of these endpoints plus a randomly selected control group were included, and all together 9528 subjects were genotyped [22]. For the present study, subjects who met to the second visit of the fourth survey and who were not genotyped in 2012 (n = 2369), were now included and genotyped for the same SNPs as the initial cohort. In addition, all subjects were also genotyped for several new SNPs.

### Measurements

At the survey in 1994–1995, the participants filled in questionnaires on medical history and lifestyle factors. Blood pressure, height and weight, serum total cholesterol and triglycerides were measured and analyzed as previously described [22]. Sera from the second visit were stored at −70°C, and after a median storage time of 13 years, thawed in March 2008 and analyzed for 25(OH)D using an automated clinical chemistry analyzer (Modular E170, Roche Diagnostics®, Mannheim, Germany). The assay overestimates the serum 25(OH)D levels in smokers [24], which was corrected for in the statistical analyses.

### Selection of SNPs

Instead of analyzing all SNPs related to the serum 25(OH)D level separately, we created a genotype score based on two 25(OH)D synthesis SNPs (rs12785878 in the *NADSYN1* gen responsible for the availability of 7-dehydrocholesterol in the skin and rs10741657 in the *CYP2R1* gene involved in the conversion of vitamin D into 25(OH)D) that in our cohort were the ones most strongly related to the serum 25(OH)D level. One point was given for the homozygote genotype with highest serum 25(OH)D, two for the heterozygote, and three for the homozygote with the lowest level, and the points for the two SNPs added together. Since SNPs in the *VDR* have been less studied in relation to mortality, we included the four most commonly analyzed *VDR* SNPs (rs7975232 (Apa1), rs1544410 (Bsm1), rs2228570 (Fok1), and rs731236 (Taq1)). In addition, we also included two *VDR* SNPs (rs2239179 and rs7968585) and two *CUBILIN* SNPs (rs1801222 and rs12766939) that recently were reported to have an interaction with the serum 25(OH)D regarding a composite clinical outcome [25]. A *MEGALIN* SNP (rs3755166) was included since the transport of the vitamin D-DBP complex into the renal tubuli cells depends on the endocytic cubulin/megalin system [26].

### Genotyping

Genotyping of these SNPs was performed in blood samples collected in the fourth survey of the Tromsø study in 1994-1995 by KBiosciences (http://www.lgcgenomics.com/genotyping/) using a competitive allele-specific polymerase chain reaction (KASPar) assay that enables highly accurate bi-allelic scoring of SNPs, as previously described in detail [22].

### Statistical analyses

The relations between SNP genotypes and mortality were evaluated in Cox regression analyses with age and gender as covariates and with the major homozygote genotype used as reference. The observation time was set from 1994-1995, and the period of observation was cut off by March 2017. Information on death was obtained from the Causes of Death Registry, updated till March 2017. In further analyses, risk factors for mortality as BMI, systolic blood pressure, serum lipids, and smoking were included as covariates to examine if relations could be explained through these risk factors.

Distribution of the continuous variables serum 25(OH)D, blood pressure, lipids and BMI was evaluated for skewness and curtosis and visual inspection of histograms and found normal except for serum triglycerides which was normalized by log transformation before use as dependent variable. Trends across the genotypes were evaluated with linear regression with age and gender as covariates.

For the relation between serum 25(OH)D and mortality, adjustments for season and smoking were performed by calculating 25(OH)D percentiles for each month for smokers and non-smoker separately. Based on this, the cohort was then divided in the following serum 25(OH)D percentile groups: 0-10 percentile, 11-25 percentile, 26-50 percentile, and > 50 percentile.

The genotype frequencies were examined for compliance with Hardy-Weinberg equilibrium using chi-squared analysis [27]. Linkage disequilibrium (LD) between SNPs was evaluated with r^2^ and Lewontin’s D′ statistics [28,29].

The data are shown as mean ± SD. All tests were done two-sided, and a P-value < 0.05 was considered statically significant. The P-values are shown without corrections for multiple comparisons.

### Ethics

The study was approved by the Regional Committee for Medical and Health Research Ethics (REK Nord) (reference 2010/2913-4). Only participants with valid written consent were included.

## Results

### Baseline

A total of 11 897 subjects had at least one SNP successfully analyzed, 5347 (44.9%) men and 6550 (55.1%) women, and were included in the analyses. Among these, 33.7 % were current smokers at baseline. In 1994-1995 their mean ± SD age was 57.8 ± 13.6 years and BMI 25.9 ± 4.1 kg/m^2^.

The individual SNP analyses were successful in 98.8–99.5% of the subjects. The genotypes for all the SNPs were in Hardy-Weinberg equilibrium except for rs12766939 in the *CUBILIN* gene (Chi square 8.36, P < 0.01), and rs12785878 in the *NADSYN1* gene (Chi square 7.97, P < 0.01). The *VDR* SNPs rs7975232 (Apa1), rs1544410 (Bsm1), and rs731236 (Taq1) were in LD with each other (r^2^ > 0.4) and also with rs2239179 and rs7968585 (r^2^ > 0.4). Rs2239179 and rs7968585 were in moderate LD (r^2^ = 0.386). None of the other SNPs (including rs2228570 (Fok1)) were in LD.

Among the 11 897 subjects, serum 25(OH)D was analyzed in 6733 subjects. The difference in serum 25(OH)D between the major and minor homozygote for the two selected SNPs in the *CYP2R1* and *NADSYN1* genes were 2.8 and 2.0 nmol/L, respectively. For the other SNPs tested, there was only a significant difference of 1.5 nmol/L in serum 25(OH)D for the *VDR* SNP rs2228570 (Fok1). The difference in serum 25(OH)D between highest and lowest genotype score was 3.6 nmoL/L (Table 1).

**Table 1.**
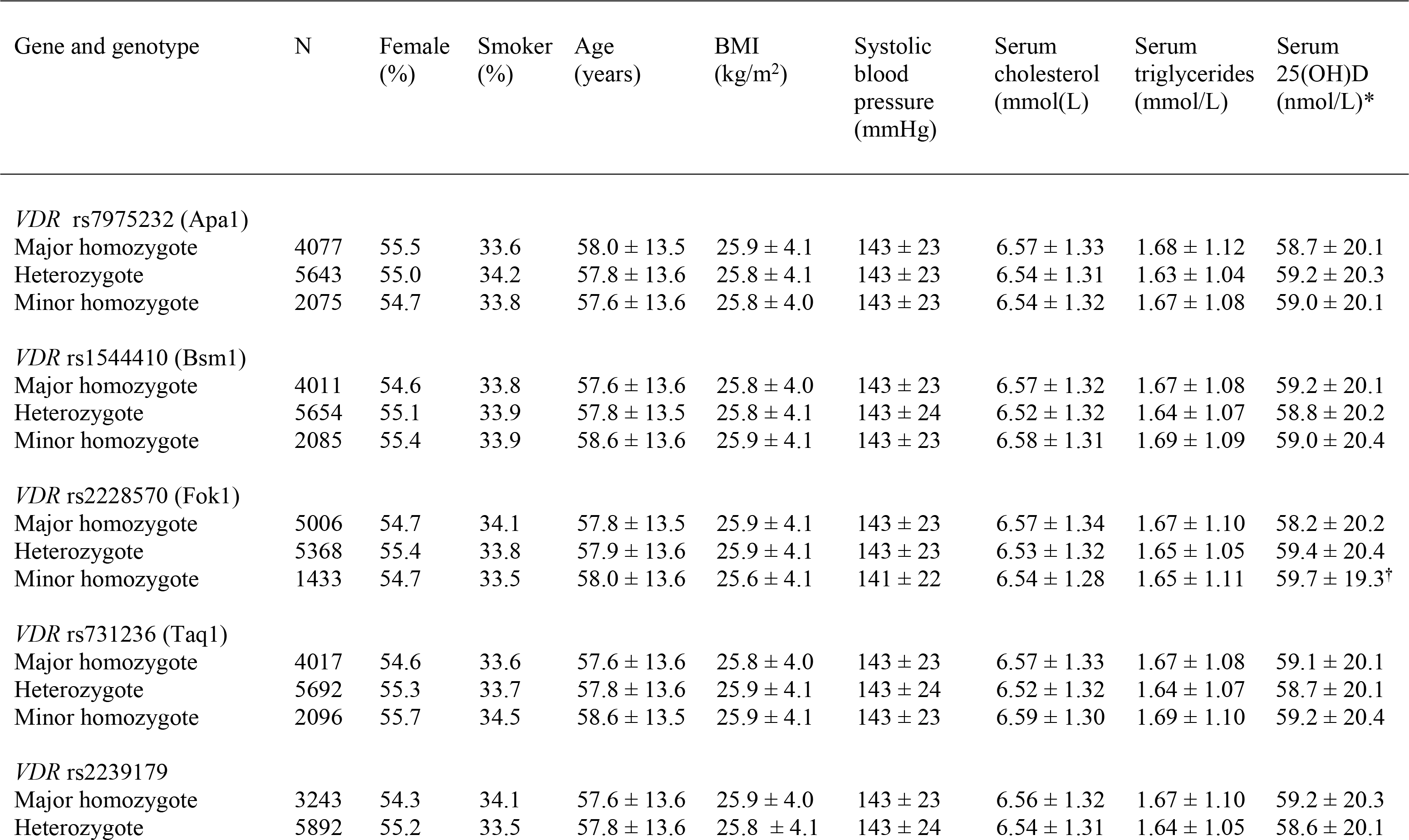
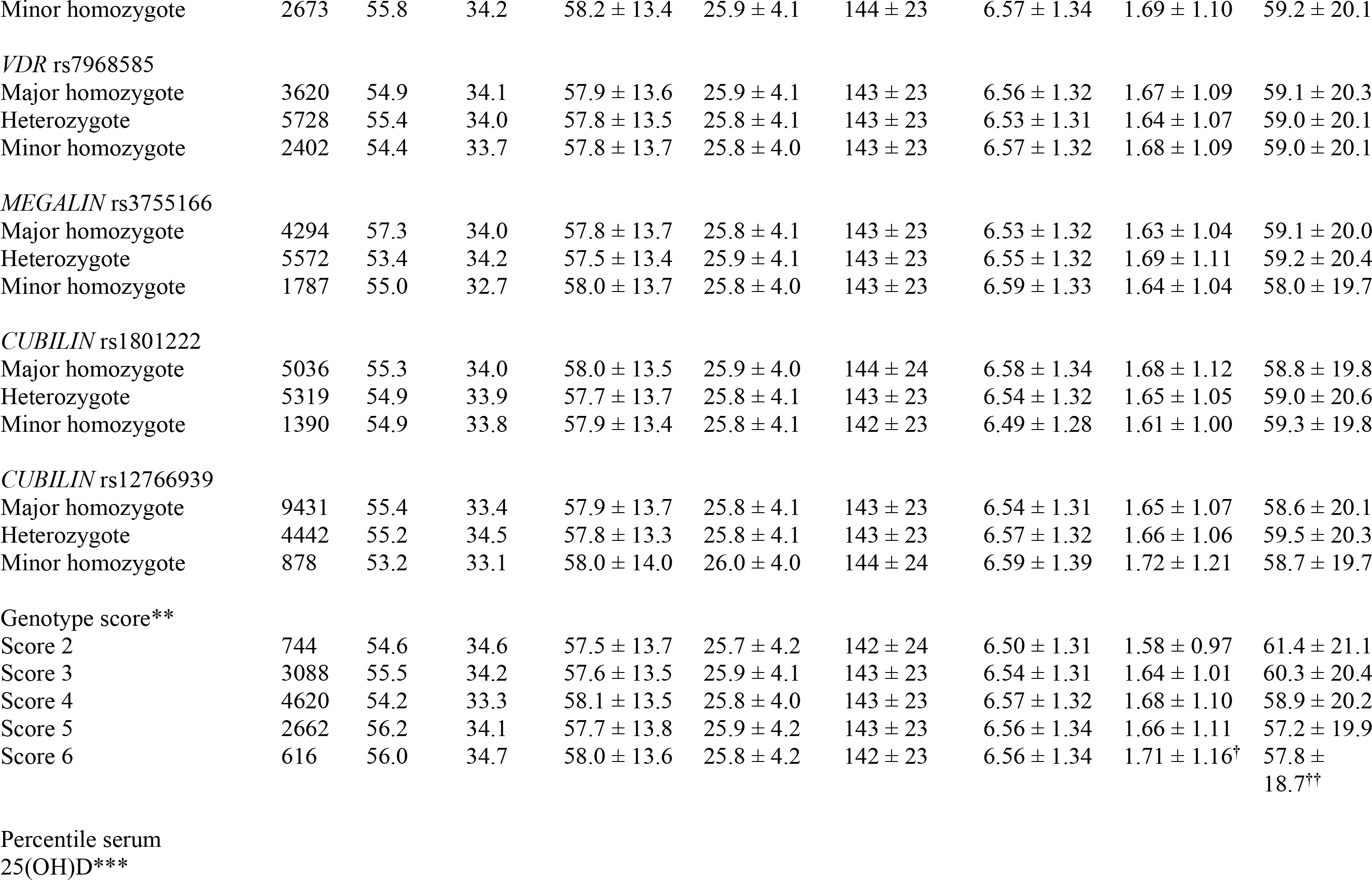
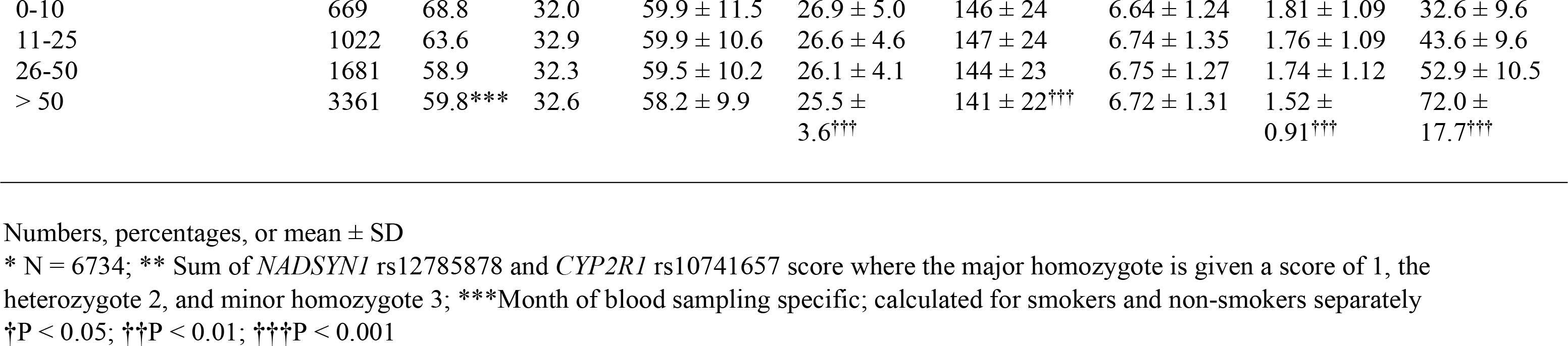
Baseline characteristics (1994-1995) in relation to genotypes. The Tromsø Study

There were no statistically significant differences between the genotypes regarding age and gender, smoking status, systolic blood pressure or serum cholesterol. However, for the genetic score, there was a significant association with serum triglycerides with the highest triglyceride levels in those with the highest genetic score (lowest serum 25(OH)D) (linear trend, P = 0.023).

Subjects in the lowest serum 25(OH)D percentile group had higher BMI, systolic blood pressure and serum triglyceride levels than those in higher serum 25(OH)D percentiles (linear trend, P < 0.001) (Table 1).

### Cox regression

During the observation period 5491 subjects (2762 men and 2729 women) had died. In the Cox regression analysis, only the *CUBILIN* SNP rs12766939, the genotype SNP score, and the serum 25(OH)D percentile groups were significantly associated with mortality (Table 2) and Figures 1–3. These associations were not significantly affected by inclusion of systolic blood pressure, BMI, smoking status, serum cholesterol and serum triglycerides in the Cox regression analysis. When including serum 25(OH)D as a continuous variable (reducing the number of subjects to 6655), the same patterns for *CUBILIN* SNP rs12766939 and the genotype SNP score and mortality were seen, but the relations were no longer statistically significant (data not shown). There was no significant interaction between the *CUBILIN* SNP rs12766939 (or any of the other SNPs) and the serum 25(OH)D level regarding mortality.

**Figure 1.**
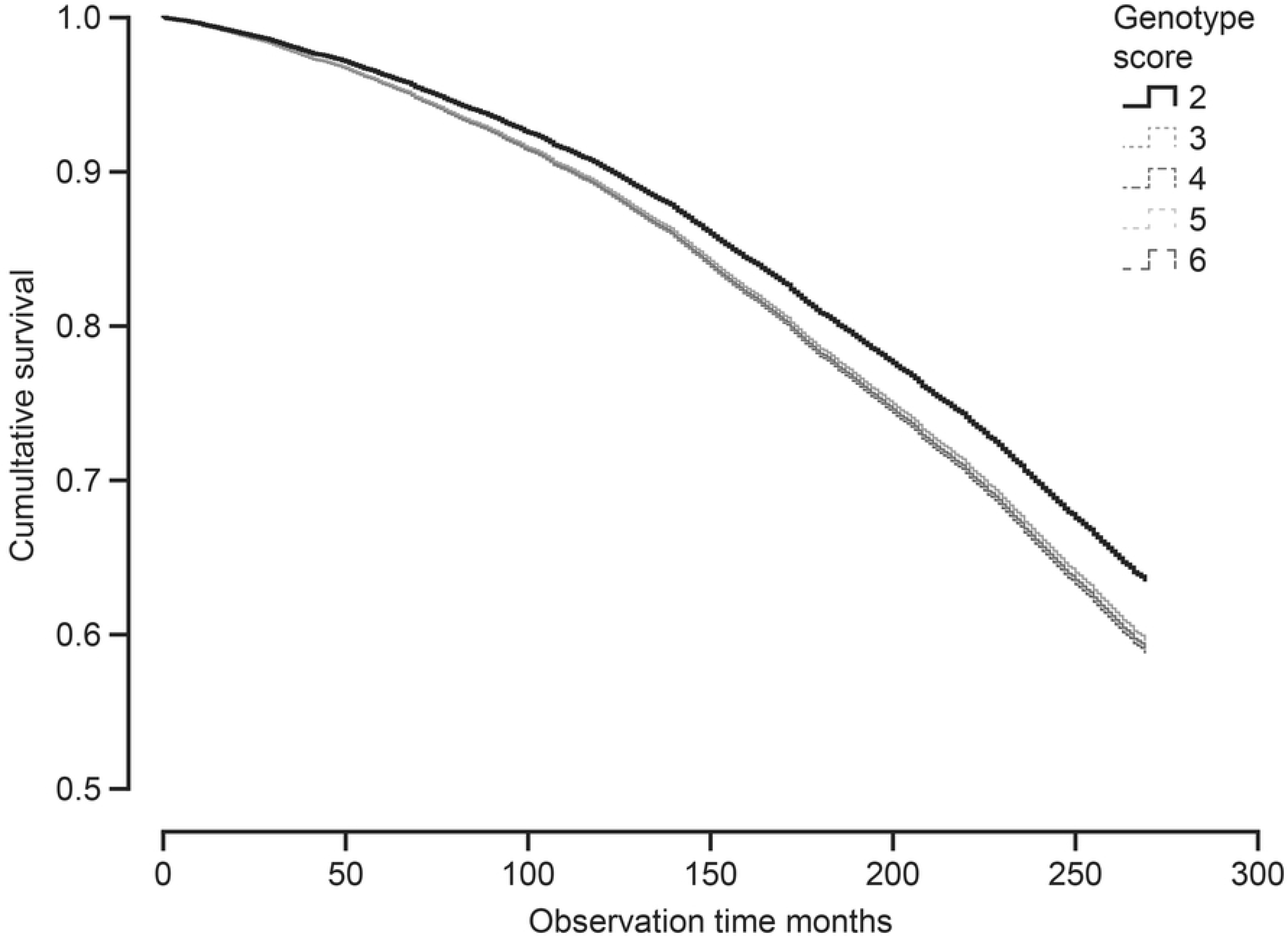
Cumulative probability of survival according to genotype score based on SNP rs12785878 in the *NADSYN1* gene and SNP rs10741657 in the *CYP2R1* gene, Cox regression with age and sex as covariates. Low genotype score corresponds to high serum 25(OH)D level.

**Figure 2.**
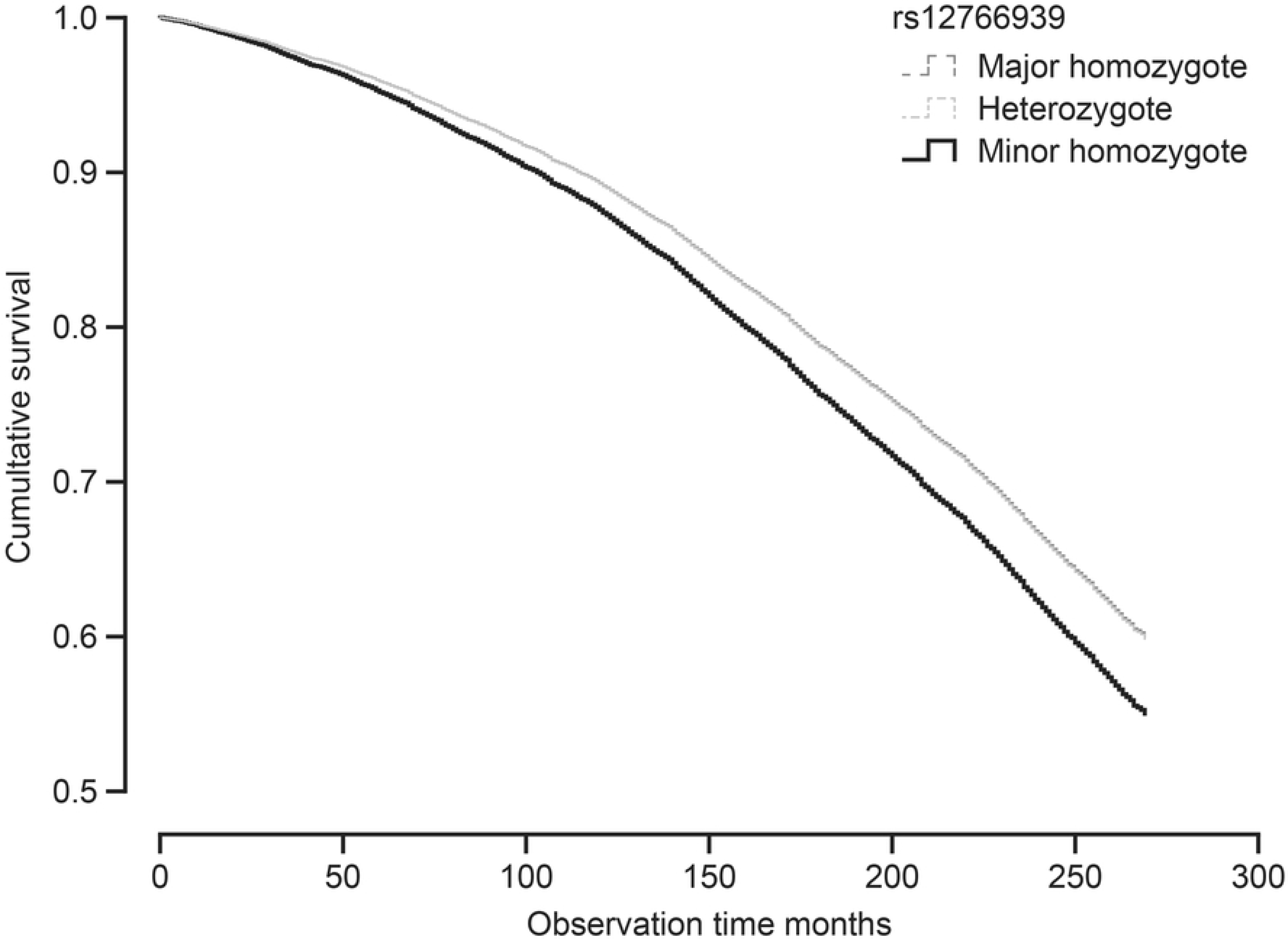
Cumulative probability of survival according to rs12766939 genotypes (in the *CUBILIN* gene), Cox regression with age and sex as covariates.

**Figure 3.**
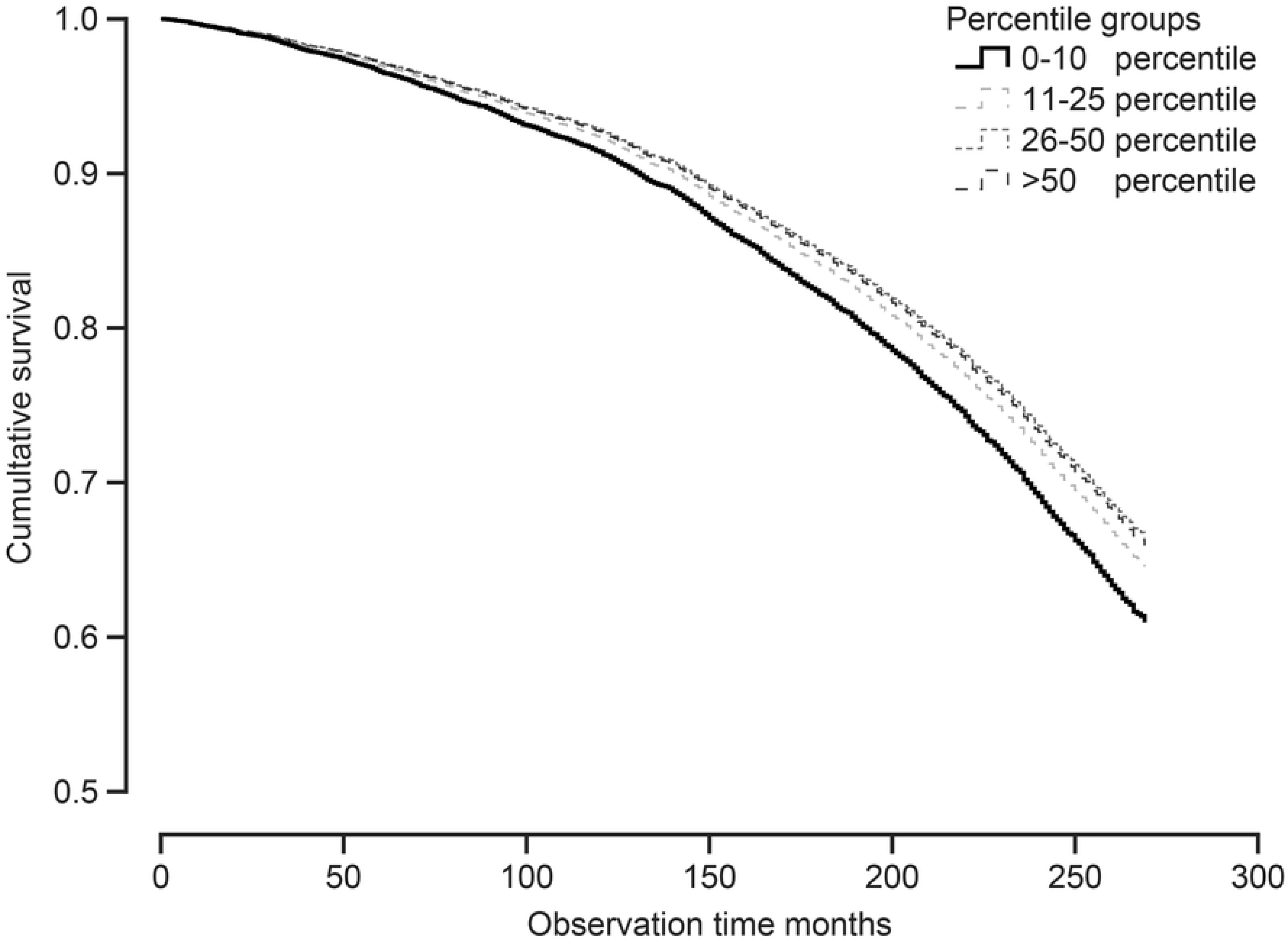
Cumulative probability of survival according to serum 25(OH)D percentile score group (month specific), Cox regression with age and sex as covariates.

**Table 2.** Hazard ratios of death in relation to genotypes, genotype score, and serum 25(OH)D percentile groups. The Tromsø Study 1994-2015.

| Gene and genotype | N | Dead (%) | Hazard ratio | 95 % confidence interval | Significant P values |
| --- | --- | --- | --- | --- | --- |
| *VDR* rs7975232 (Apa1) |  |  |  |  |  |
| Major homozygote | 4077 | 46.6 |  | Reference |  |
| Heterozygote | 5643 | 45.5 | 0.982 | 0.926 – 1.042 |  |
| Minor homozygote | 2075 | 47.0 | 0.996 | 0.922 – 1.076 |  |
| *VDR* rs1544410 (Bsm1) |  |  |  |  |  |
| Major homozygote | 4011 | 47.2 |  | Reference |  |
| Heterozygote | 5654 | 44.8 | 0.963 | 0.907 – 1.022 |  |
| Minor homozygote | 2085 | 48.4 | 1.018 | 0.943 – 1.099 |  |
| *VDR* rs2228570 (Fok1) |  |  |  |  |  |
| Major homozygote | 5006 | 6.0 |  | Reference |  |
| Heterozygote | 5368 | 46.2 | 0.993 | 0.938 – 1.051 |  |
| Minor homozygote | 1433 | 56.6 | 1.015 | 0.931 – 1.106 |  |
| *VDR* rs731236 (Taq1) |  |  |  |  |  |
| Major homozygote | 4017 | 47.1 |  | Reference |  |
| Heterozygote | 5692 | 44.7 | 0.962 | 0.907 – 1.022 |  |
| Minor homozygote | 2096 | 48.4 | 1.021 | 0.946 – 1.102 |  |
| *VDR* rs2239179 |  |  |  |  |  |
| Major homozygote | 3243 | 46.4 |  | Reference |  |
| Heterozygote | 5892 | 45.3 | 1.003 | 0.942 – 1.069 |  |
| Minor homozygote | 2673 | 47.5 | 1.037 | 0.963 – 1.118 |  |
| *VDR* rs7968585 |  |  |  |  |  |
| Major homozygote | 3620 | 46.5 |  | Reference |  |
| Heterozygote | 5728 | 45.2 | 0.965 | 0.907 – 1.026 |  |
| Minor homozygote | 2402 | 47.7 | 1.006 | 0.933 – 1.084 |  |
| *MEGALIN* rs3755166 |  |  |  |  |  |
| Major homozygote | 4294 | 46.6 |  | Reference |  |
| Heterozygote | 5572 | 46.0 | 1.025 | 0.966 – 1.086 |  |
| Minor homozygote | 1787 | 44.8 | 0.943 | 0.869 – 1.024 |  |
| *CUBILIN* rs1801222 |  |  |  |  |  |
| Major homozygote | 5036 | 47.1 |  | Reference |  |
| Heterozygote | 5319 | 45.7 | 0.966 | 0.912 – 1.022 |  |
| Minor homozygote | 1390 | 44.3 | 0.935 | 0.856 – 1.022 |  |
| *CUBILIN* rs12766939 |  |  |  |  |  |
| Major homozygote | 9431 | 45.9 |  | Reference |  |
| Heterozygote | 4442 | 45.5 | 1.005 | 0.949 – 1.063 |  |
| Minor homozygote | 878 | 50.8 | 1.172 | 1.061 – 1.295 | 0.002 |
| Genotype score* |  |  |  |  |  |
| Score 2 (high 25(OH)D) | 744 | 41.1 |  | Reference |  |
| Score 3 | 3088 | 46.3 | 1.167 | 1.032 – 1.321 | 0.014 |
| Score 4 | 4620 | 47.1 | 1.159 | 1.028 – 1.307 | 0.016 |
| Score 5 | 2662 | 45.6 | 1.137 | 1.003 – 1.289 | 0.044 |
| Score 6 | 616 | 46.8 | 1.156 | 0.984 – 1.358 |  |
| Percentile serum 25(OH)D** |  |  |  |  |  |
| 0-10 | 669 | 48.0 | 1.188 | 1.051 – 1.344 | 0.006 |
| 11-25 | 1022 | 46.3 | 1.050 | 0.945 – 1.167 |  |
| 26-50 | 1681 | 42.8 | 0.981 | 0.897 – 1.074 |  |
| > 50 (high 25(OH)D) | 3361 | 40.0 |  | Reference |  |

Cox regression with adjustment for age and gender
\* Sum of *NADSYN1* rs12785878 and *CYP2R1* rs10741657 score where the major homozygote is given a score of 1, the heterozygote 2, and minor homozygote 3; \*\*Month of blood sampling specific; calculated for smokers and non-smokers separately

## Discussion

In the present study we have confirmed the relation between low serum 25(OH)D and mortality. We have also found a weak relation between genotype score (based on two SNPs related to the 25(OH)D level) and mortality, and possibly also a relation between one SNP in the *CUBILIN* gene and mortality. However, no significant relations between *VDR* and *MEGALIN* SNPs and mortality were found.

As for the relation between serum 25(OH)D and mortality, this is well established in numerous studies [30–33], and was in our study not affected by adjusting for sex or age, nor for the cardiovascular risk factor smoking, BMI, systolic blood pressure, or serum lipids. Due to the high risk of reverse causality, as a low serum 25(OH)D may be the result as well as the cause of disease, associations have to be confirmed in properly performed RCTs. However, there has been no RCT performed specifically for vitamin D and mortality, but meta-analyses of available data have indicated a protective effect by vitamin D supplementation [15,34]. The validity of this conclusion has, however, been questioned [35].

An approach to eliminate effects of confounders is MR studies, but so far there is only one large MR study on vitamin D and mortality. Thus, Afzal et al. included 95 766 subjects from three Danish cohorts, and during mean observation times of 5 to 19 years, 10 349 subjects died. SNPs in the *DHCR7* (*NADSYN1*) and *CYP2R1* genes were used to create a genotype score similar ours, and a genetically 20 nmol/L lower serum 25(OH)D level gave an odds ratio for mortality of 1.30 (1.05 – 1.61) [17]. Even if this study is impressive, it should be recalled that this group also published a significant association between 25(OH)D SNPs and type 2 diabetes in a cohort of 96 423 subjects with 5037 cases [36], a finding that could not be reproduced in an even larger study [11].

It was therefore prudent that the editorial following the Afzal et al. mortality publication asked for confirmatory studies [18], and several more, but smaller, MR studies have now been published. Thus, Ordonez-Mena et al. included 8417 subjects of whom 1338 died during the mean observation time of 11 years, but found no relation to serum 25(OH)D associated SNPs [37]. On the other hand, in the study by Aspelund et al. [38] who included 10 501 subjects from three European cohorts (including parts of our present cohort) where 4003 subjects died during a median observation time of 10.4 years, the relation between a 25(OH)D genotype score and mortality was comparable to the one reported by Afzal et al. [17]. However, it did not reach statistical significance, probably due to lack of power. This is very similar to our findings, with a ~ 15 % increased mortality risk in those with less favourable genotype scores (associated with lower serum 25(OH)D) compared to the ones with the most favourable score. However, this would not reach statistical significance if adjusted for multiple testing. Furthermore, rs12785878 in the *NADSYN1* gene, which was part of our genotype score, was not in Hardy-Weinberg equilibrium. This can be explained by the high proportion with Sami ancestry in the Tromsø population [39], as this SNP has a considerable difference in allele frequency between Asian and Western populations [40]. Our results could therefore be biased, and the findings by Afzal et al. [17] on mortality and vitamin D are so far not confirmed with certainty.

For 1,25(OH)_2_D, the active form of vitamin D, to have an effect, it has to connect to its nuclear receptor VDR. Mutations in the *VDR* could change the affinity of the receptor for 1,25(OH)_2_D, and numerous *VDR* SNPs have been described. However, the biological functions of these SNPs are uncertain, and effects associated with them ascribed to haplotype connections or linkage to truly functional polymorphisms elsewhere in the VDR [41]. This is the case for the most studied *VDR* SNPs Apa1, Bsm1, Taq1, and Fok1 that in particular have been analyzed in relation to cancer, but with inconsistent results [12]. There are only a few studies on these *VDR* SNPs and mortality, and as for cancer the results are non-conclusive. Thus, de Jongh et al. included 923 subjects of whom 480 participants deceased during the median follow up time of 10.7 years, but found no significant relations to mortality for the Apa1, Bsm1, Taq1, and Fok1 SNPs, nor when analyzing their haplotypes [19]. Similarly, Perna et al. found no relation between *VDR* polymorphisms and mortality in 1397 colorectal cancer patients [20], whereas Marco et al. found a relation between Bsm1 polymorphism and survival in 143 subjects on hemodialysis [21]. To our knowledge our study is therefore the largest where these SNPs have been analyzed in relation to mortality, but the results were negative.

We also included two other *VDR* SNPs (rs2239179 and rs7968585) since they in a study by Levin et al. were found to modify the association between serum 25(OH)D and a composite outcome consisting of hip fracture, myocardial infarction, cancer and mortality [25]. However, we could not find an association between these SNPs and mortality, nor did we find an interaction with low serum 25(OH)D levels. This is in agreement with the studies by Ordonez-Mena et al. [37] and Vimaleswaran et al. [42]. Even though the concept of interaction between an altered VDR receptor and vitamin D deficiency is appealing, this still needs supportive data.

Other elements in the vitamin D metabolism are the endocytic receptors megalin and cubulin, which are present in the renal tubuli cells [43] and enable transportation of the DBP-vitamin D complexes and other filtered proteins into the cells [26]. Cubilin dysfunction may lead to urinary loss of vitamin D in the urine and cause abnormal metabolism of 25(OH)D [44], and *MEGALIN* polymorphisms are associated with adiposity [45]. Furthermore, an interaction between *CUBILIN* SNPs and serum 25(OH)D level was also described by Levin et al. [25]. Again, we could not confirm such interactions, but the *CUBILIN* SNP rs12766939 was significantly associated with mortality, which to our knowledge has not been reported before. It should be noted though that this SNP, similar to rs12785878 in the *NADSYN1* gene, was not in Hardy-Weinberg equilibrium and probably for the same mixed-population reason. Our results should therefore be considered as exploratory and need confirmation in other populations. Furthermore, this SNP was not associated with lower serum 25(OH)D levels, and the association to mortality probably more related to altered reabsorption of other proteins or protein-complexes in the renal proximal tubule or the intestine [46].

Our study has several important limitations. Although there is a fairly strong heritability of the serum 25(OH)D level, which from twin studies has been estimated to be between 20 to 85 % [47], our genotype score can only explain a fraction of this heritability. Recently a SNP in the *CYP2R1* gene was identified with a 3 to 4 fold larger effect size on the serum 25(OH)D level than the *CYP2R1* SNP included by us [8]. This SNP has a minor allele frequency < 5% and shows a strong relation to risk of multiple sclerosis [8]. Genotyping for this SNP (and similar SNPs detected when SNPs with low minor allele frequency related to serum 25(OH)D are sought for) would of course give the MR analysis much more power. Furthermore, our *VDR* SNPs are probably non-functional, and significant clinical effects unlikely unless these SNPs by chance are coupled to truly functional SNPs. We did not adjust for multiple testing, which would have made the relation with mortality for the genotype score and the serum 25(OH)D percentile groups non-significant. Some of our SNPs were not in Hardy Weinberg equilibrium. The results should therefore be interpreted with caution, and the relation between the *CUBILIN* SNP rs12766939 and mortality (which was highly significant even after Bonferroni correction) should be considered as exploratory.

Our study also has strengths. It is, to our knowledge, the largest on *VDR*, *CUBILIN* and *MEGALIN* SNPs and mortality, and the results regarding genotype score are in line with similar studies.

In conclusion, the questions regarding vitamin D and mortality are still unanswered. Although MR studies also have their limitations [5], discovery of SNPs more strongly related to serum 25(OH)D levels and SNPs with direct effects on the function of the VDR, might hopefully bring us closer to an answer.

## Acknowledgements

The study was supported by grants from the North Norway Regional Health Authorities; The Norwegian diabetes Association; The research council of Norway and UiT The Arctic University of Norway

## Author contributions

Conceptualization, R.J. and G.G.; Methodology, R.J., T.W. and G.G.; Formal Analysis, R.J. and G.G.; Resources, R.J., T.W. and G.G; Data Curation, T.W.; Writing – Original Draft Preparation, R.J.; Writing – Review & Editing, R.J., T.W. and G.G.; Project Administration, T.W.; Funding Acquisition, R.J.and G.G.”.

